# Functional phenomics: An emerging field integrating high-throughput phenotyping, physiology, and bioinformatics

**DOI:** 10.1101/370288

**Authors:** Larry M. York

## Abstract

**Highlight:** Functional phenomics is an emerging field in plant biology that relies on high-throughput phenotyping, big data analytics, controlled manipulative experiments, and simulation modelling to increase understanding of plant physiology.

**Abstract:** The emergence of functional phenomics signifies the rebirth of physiology as a 21st century science through the use of advanced sensing technologies and big data analytics. Functional phenomics highlights the importance of phenotyping beyond only identifying genetic regions because a significant knowledge gap remains in understanding which plant properties will influence ecosystem services beneficial to human welfare. Here, a general approach for the theory and practice of functional phenomics is outlined including exploring the phene concept as a unit of phenotype. The functional phenomics pipeline is proposed as a general method for conceptualizing, measuring, and validating utility of plant phenes. The functional phenomics pipeline begins with ideotype development. Second, a phenotyping platform is developed to maximize the throughput of phene measurements. A mapping population is screened measuring target phenes and indicators of plant performance such as yield and nutrient composition. Traditional forward genetics allows genetic mapping, while functional phenomics links phenes to plant performance. Based on these data, genotypes with contrasting phenotypes can be selected for smaller yet more intensive experiments to understand phene-environment interactions in depth. Simulation modeling can be used to understand the phenotypes and all stages of the pipeline feed back to ideotype and phenotyping platform development.

## Introduction

Global food insecurity is among the most significant challenges for the 21st century (Grafton *et al.*, 2015). Arguably, advances in genetics and genomics have far out-paced understanding of plant physiology. However, for plant science to become positioned to offer solutions to agriculture’s great challenges, understanding the fundamental processes contributing to crop performance must be prioritized. High-throughput phenotyping has allowed unprecedented data collection on plant form and function (Pieruschka and Poorter, 2012), yet its utility is commonly framed in the context of gene discovery. Phenomics, research focused on understanding variation in plant phenotypes (Furbank and Tester, 2011), has quickly become a biological discipline, yet is often secondary to a focus on genetics. Functional phenomics is proposed as a field of inquiry that leverages high-throughput phenotyping to create knowledge about plant function at the physiological level. The value of the knowledge generated by functional phenomics justifies the endeavour, not necessarily requiring the simultaneous study of the underlying genetics.

Phenomics itself is a relatively young field of plant science which emerged by the early 2000s (Edwards and Batley, 2004; Holtorf *et al.*, 2002), but previously established in mouse research (Gerlai, 2002) and microbiology (Schilling *et al.*, 1999). Broadly, the literature has consistently used phenomics in the context of a field of research focused on understanding the diversity of plant phenotypes, and usually with special consideration of high-throughput phenotyping (Furbank and Tester, 2011; Pieruschka and Poorter, 2012). Phenomics research often measures diverse species or genotypes in order to describe the variation of target phenes (fundamental units of phenotype of replacing the world “trait,” see Box 1) and to link these measures to the underlying genetics (Atkinson *et al.*, 2015). Therefore, substantial knowledge has been created about phene-gene maps, yet relatively less is known about the phene-function map. In contrast, functional phenomics is concerned primarily with understanding the relationship of phene variation to functional variation. The functional phenomics pipeline offers an avenue for designing and implementing research leveraging high-throughput phenotyping and data analytics.

## The Functional Phenomics Pipeline

### Ideotype development

The first stage of the pipeline is ideotype development (Box 2). An ideotype is the ideal phenotype in an environment to either maximize capital gains (or yield) in agriculture (Donald, 1968), or to maximize fitness in natural systems. Donald’s (1968) wheat ideotype was proposed as being short, having erect leaves, and an erect, large ear, among other properties which are still influencing wheat breeding to this day. Shoot architecture and growth determinism have been ideotype targets in beans (Kelly, 2001). In order to optimize root systems in maize, Lynch (2013)proposed the steep, deep, and cheap ideotype to enhance efficient soil exploration and exploitation. In all cases, the determination of ideotype was environment-dependent, reflecting an inherent ecophysiological approach. Conveniently, not only is the concept of ideotype useful for plant breeding, but also for framing our understanding of plant function. In that regard, the ideotype serves as a set of hypotheses about how plants work. In the conception of an ideotype, what phenes are likely to influence performance is a central question, so ideotype development is partially synonymous with phene determination. A crucial premise for the utility of functional phenomics in plant breeding is that ideotype-based breeding will eventually generate more yield gains than selection for yield alone. The rationale for this prediction is that selection for yield alone, given finite population sizes with limited genetic and phenetic shuffling, cannot jump across local depressions in the yield-phene performance landscape (Messina *et al.*, 2011) to reach even higher peaks, or that all beneficial phene states are unlikely to all be present in a single genotype. In contrast, ideotype breeding supposes that breeders can be plant engineers that design the optimal plants by stacking beneficial phene states.

### Phenotyping platform

Once ideotype development has led to the set of phenes hypothesized to most greatly influence crop performance, a phenotyping platform can be designed to maximize throughput for determination of the proposed phenes’ states across many samples. Among many considerations for a phenotyping platform, a primary one is the tradeoff between extensive (many samples) measurements and intensive (many measurements) phenotyping (Fiorani and Schurr, 2013). The more manual requirements of a platform, the stronger this tradeoff becomes. Fortunately, developments in technology alleviate this tradeoff by allowing many precise measurements to be made quickly for each sample. Broadly, these developments are in electronic sensors, computational power, and available software. Specifically, image-based phenotyping has quickly become the dominant mode using readily available digital cameras and automated image analysis software for determining phenes related to plant architecture and light reflectance profiles (Minervini *et al.*, 2015).

Perhaps it is useful to consider the full phenotyping platform as a fusion of a growth platform and a measurement platform. How to grow the plants affects throughput, but more importantly affects the actual structure and function of the plant. Lab and greenhouse-based screens are usually higher throughput, yet in some situations may not correlate well to field-based measures (Poorter *et al.*, 2016). However, even year-to-year correlations can be low in the field so the relative usefulness of all growth platforms is still an open question.

Another aspect is how to bring the sensor in proximity to the plant, with two possibilities: plant-to-sensor or sensor-to-plant, as utilized by unmanned aerial systems, robots, over-canopy carts or tractors (Pittman *et al.*, 2015), and in some greenhouse systems (Vadez *et al.*, 2015). Sensor-to-plant technology is especially relevant for field-based phenotyping as removing the plant is always destructive, but in the greenhouse plants can be grown in pots and moved by conveyor or other means to the sensors (Campbell *et al.*, 2015). Plant-to-sensor is also commonly used in destructive root phenotyping in the field, such as root crown phenotyping (York, 2018), or shovelomics, as well as for spectroradiometry of plant leaves (Biliouris *et al.*, 2007).

Functional phenomics seeks to relate phenes to plant performance, or fitness. Performance metrics could be classified as phenes, yet at the same time they are clearly the outcomes of other aspects of plant phenotype. Therefore, functional phenomics extends more common phenotyping research to not only include the phenes of interest but also measures of plant performance. The diversity of phenes measured may necessitate the use of several phenotyping platforms. More traditional and manual plant performance metrics include shoot dry mass, leaf area, and nutrient content through manual weighing, scanning, and chemical analysis, respectively. However, image-based shoot phenotyping can supplant these methods using automated measures of plant volume, height, leaf area, and spectral indices that reflect plant nutrient and water status (Haghighattalab *et al.*, 2016; Hunt *et al.*, 2013). Advances in high-throughput measures of plant transpiration through use of infrared imagery (Deery *et al.*, 2014) and photosynthesis through use of chlorophyll fluorescence and rapid gas-exchange measurements (Stinziano *et al.*, 2017) will be especially important over the next decade.

### Phenotyping a population

A central premise of the functional phenomics pipeline is that variation in measured phenes can be linked to variation in functional outcomes. Therefore, use of diverse germplasm is essential in order to ensure phenotypic and functional variation. Biparental populations of recombinant inbred lines are commonly used for mapping, however the phenotypic and genetic variation is limited to the original parents. As sequencing has become cheaper, use of large diversity panels including more phenotypic and genetic variation have become more common. In both cases, recombination serves to shuffle genetic elements. The relatedness of the individuals is needed for the genetic analysis, and while known for families, the population structure of diversity panels is usually unknown so must be inferred. Errors in the inference of population structure can lead to spurious associations among phenes (Myles *et al.*, 2009) and are therefore a major problem for the use of diversity panels. Using mapping populations for functional phenomics allows mapping phenes to functions as well as genes, generating synergistic opportunities.

### Bioinformatics of functional phenomics

Arguably, the greatest challenge faced by functional phenomics is not the acquisition of mass amounts of phenotypic data, but the analysis of the data in order to create knowledge (Tardieu *et al.*, 2017). Most phenotypic data being generated is continuous and numeric, for both phenes and performance indicators. Therefore, regression techniques are suitable for relating phene to function, and as genotype numbers and replications increase, so does statistical power. Regression can be used for testing hypotheses about phene-function relationships, but at risk of being considered ‘p-hacking’, another powerful approach is for hypothesis generation about previously unknown relations. Such data mining and testing of multiple statistical models will undoubtedly be commonplace in functional phenomics, and therefore needs to be reported in methods sections of manuscripts to avoid inflation bias (Head *et al.*, 2015). At the same time, the value of data mining for hypothesis generation must be acknowledged and not stigmatized.

Correlational analysis of the phene network (Box 3) is an important aspect of functional phenomics. Pairwise correlations of all measured phenes can identify collinearity among phenes, which is important for downstream decisions about statistics. Phene correlations also raise the question: why? Do phenes correlate because of a shared developmental program? Do they serve a similar purpose? Such questions are implicit in the concept of phene modules (Murren, 2002), defined as correlated phenes acting within a functional unit (such as a flower). Phene A may be correlated to phene B for four reasons: A partially controls B, B partially controls A, another phene C partially controls both A and B, or the correlation is simply spurious and not repeatable. These statistical frameworks may also identify patterns in phene hierarchies, possibly defining which phenes are the major drivers, and which might be considered phene aggregates comprised of lower order phenes (York *et al.*, 2013). Phenetic hierarchies may be determined through the use of structural equation modelling (Tetard-Jones *et al.*, 2014) and network analysis (Bartsch *et al.*, 2015).

Principal-component analysis yields scores that are the linear combinations of underlying phenes that maximize the explained multivariate variation. The loadings that multiply the individual phenes to sum to the component and give an indication of which phenes are most strongly associated with a principal component. Indeed, the entire component can be viewed as a latent variable that drives the variation of the constituent phenes. These latent variables can be described as ‘dark’ phenes that influence measured phenes without being readily measured themselves, and are related to the concept of cryptotype (Chitwood and Topp, 2015). In recent works, such dark phenes of root system architecture were mapped to genetic regions in soybean (Dhanapal *et al.*, 2018). In wheat dark phenes where inferred based on measures of internal consistency among shoots and tillers for the same phenes and principal components were used to construct estimations of the latent constructs (York *et al.*, 2018).

The methods for linking phene variation to performance metrics are similar as for the phene correlational statistics described above. Given the continuous nature of most phenes and performance metrics, regression techniques are an obvious choice for modelling the relation of phene to function or performance. Multiple regression allows several phenes to be fit to a performance measure simultaneously, and can add predictive power. Regression techniques allow mechanistic insights. For example, multiple regression of root crown phenes to shoot mass indicated that phenes related to lateral branching density, number, and angles could explain up to 68% of the variation of shoot mass, with similar results for nitrogen uptake (York and Lynch, 2015), while the maximum variation explained by a single phene was 37%. The more phenes and performance measures that are being analysed, the more observations are needed to complete the statistics. Given the multivariate nature of both phenes and performance metrics, canonical correlational analysis is another promising tool as it allows modelling of both simultaneously with multiple predictors and response variables (Gonzalez *et al.*, 2009). A fuller application of multivariate statistics is necessary to unlock the power of the functional phenomics approach.

### Confirmation of phene utility with experimentation and simulation

Phenes revealed to have relationships to crop performance must be validated before physiologists may recommend their use in breeding programs, even if the relations were previously hypothesized. Selecting genotypes that either contrast or express continuous variation for specific phenes has been a valuable approach for physiological studies that confirm phene function (Lynch, 2013; York *et al.*, 2013). While diversity panels assembled from varieties from around the world are extremely important for identifying allelic variation that influences particular phenes, diversity panels are also diverse for many phenes simultaneously. In contrast, crossing plants that contrast for a phene of interest generates a biparental population consisting of lines that often express greater variation in the phene than the parents because of transgressive segregation. Advances in gene editing technology, such as CRISPR-CAS9/Cpf1, may allow plants with contrasting phene states to be created more quickly (Gao, 2018).

Near-isophenic lines can be grown in controlled conditions for detailed physiology studies or in the field for validation under agricultural conditions. In any environment, more complete knowledge of the environment facilitates understanding of the physiological processes so the environment itself should be measured. In these experiments, the common practice is to apply abiotic stress as a treatment, such as temperature extremes, drought, or low available nitrogen, in factorial combinations with contrasting near-isophenic lines. Interpretation is simpler in cases when all plants behave similarly in the non-stressed conditions but show variation in stress conditions, otherwise differences among plants may be more related to general plant vigor rather than the phenes of interest. These controlled experiments may allow more detailed treatments, such as manipulation of plant structure, and also facilitate acquisition of more complicated measures, such as photosynthesis. The factorial designs described above that combine environmental treatments and contrasting lines are suitable for ANOVA, while the possibility of including more iso-phenic lines with more continuous variation is suitable for regression analysis.

Examples of moderate-throughput studies that directly link phenes to function are more common, and while they fall under the domain of functional phenomics, there is a continuing push to include more genotypes using high-throughput approaches to allow greater statistical power and to achieve genetic mapping. For example, decreased nighttime transpiration was shown to correlate to daytime canopy conductance and specific leaf area in maize using growth chambers fitted with scales (Tamang and Sadok, 2017). These moderate-throughput approaches are especially utilized in root biology, such as in linking root cortical senescence to root respiration and nutrient uptake (Schneider *et al.*, 2017), relating reduced secondary growth to increase phosphorus acquisition (Strock *et al.*, 2018), linking aerenchyma to living cortical area and then to yield in low phosphorus conditions (Galindo-Castaneda *et al.*, 2018), and in showing how dozens of root system architectural phenes integrate for nitrogen acquisition (York and Lynch, 2015). Scaling up the throughput of these methods that link phenes to function will strengthen functional phenomics greatly.

Simulation modelling is another promising approach in which theory of phene function can be made explicit and tested. Simulations require mechanistic understanding of how processes relate – what output is achieved by what input (Marshall-Colon *et al.*, 2017). In cases where understanding is limited, predictions from different simulations can be compared with observations to see which model is more accurate. Simulations also allow exploration of phene values beyond what has been observed in nature, possibly due to genetic or physiological constraints. Greater numbers of phenes and phene combinations can be compared in a simulation than in the field or greenhouse (York *et al.*, 2013). Finally, simulations can guide experimentation to validate phenes with the greatest potential, while new empirical data parameterize and refine the model, creating a positive feedback loop of knowledge generation (Wullschleger *et al.*, 1994). Physiological experiments and simulations, in turn, may lead to the creation of new crop ideotypes so the researcher may continue around the functional phenomics pipeline.

## Conclusion

Functional phenomics is concerned with how plants work, a knowledge gap that has been a limiting factor in understanding ecosystems and breeding for superior cultivars despite significant progress in understanding the underlying genetics. The end goal of functional phenomics is to generate new knowledge about the relationship of the plant phenome to plant functioning in ecosystems (see Box 4 for recent examples). In the context of crop breeding, the functional phenomics pipeline will guide decisions about generating populations and making selections. Functional phenomics is poised to combat major global challenges such as climate change, environmental degradation, and food insecurity.

### Box 1. What’s in a phene? A rose by any other name

The word ‘phene’ was used as early as 1925 (Serebrovsky, 1925), and some scholars have suggested the term was implied in Willhelm Johannsen’s work that coined the corresponding words phenotype, genotype, and gene (Johannsen, 1909). However, use of the term languished over the next century, appearing in some European agricultural (Gustafsson *et al.*, 1977) and ecological (Vidyakin, 2001) work, but not widely adopted. Interesting, phene-based semantics did made their way into genomics by the use of phenetic methods for genomic comparisons (Sokal, 1986; Thornton and DeSalle, 2000). However, with the rise of plant phenomics, there is a resurgence in the use of phene (Lynch and Brown, 2012; Paez-Garcia *et al.*, 2015; Pieruschka and Poorter, 2012; Rellán-Álvarez *et al.*, 2016; York and Lynch, 2015; York *et al.*, 2013). The success of phenomics in delivering solutions to global challenges will be partially determined by having adequate terminology to explain scientific observations (Slisko, 1997).

Phene is meant as a total replacement for the term trait, which is pre-eminently popular, yet is non-biological and ambiguously used (Violle *et al.*, 2007). Trait is used across biological levels of organization, while phenes are properties of organisms. Additionally, trait is often used to refer to both the general and the specific property, such as the root number trait or the many root number trait. Unfortunately, recent usage of the word phene is not careful enough to rid the literature of this ambiguous thinking, and so there is a pressing need to standardize terminology, or else phene-based semantics will fail to achieve more than contemporary thought. Phene as the general property and phene state as the specific attribute have been proposed as analogous to the concepts of gene and allele, respectively (York *et al.*, 2013). The genetics literature is also plagued by the ambiguity of using ‘gene’ in cases where ‘allele’ would be more appropriate.

Lynch and Brown (2012) recommended that phenes be considered elementary and unique at their level of organization, whether the organ, tissue, or cellular level. Defining phenes as only units of phenotype rather than fundamental units of phenotype frees the term considerably. One of the sharpest criticisms of the phene concept is that it may be impossible to know whether a given phene is fundamental, or can be deconstructed to more elemental phenes. In fact, this fuzziness should be embraced (York and Lobet, 2017) and the concept of phene hierarchies might be useful. For example, nodal root number has been linked to maize performance (Saengwilai *et al.*, 2014; York *et al.*, 2013), yet can be deconstructed to nodal roots per whorl and number of whorls (York and Lynch, 2015). Possibly these more elemental phenes could be further dissected, or, in contrast, could be aggregated in additional measures such as root length. Despite the fuzziness, a phene-based paradigm encourages the explicit consideration of these phenetic relations, and the process can yield new phenes that may be useful for understanding plant physiology. Future work will undoubtedly seek to offer further classification and understanding of phene hierarchies.

### Box 2. The functional phenomics pipeline

The functional phenomics pipeline outlines a series of research objectives that together lead to increased understanding of plant functioning by leveraging high-throughput phenotyping and data analytics. The functional phenomics pipeline begins with ideotype development to hypothesize the idea phenotyping for a given environment. Second, a phenotyping platform is developed to maximize the throughput of phene measurements related to that ideotype. Then, a mapping population is screened measuring target phenes and indicators of plant performance such as yield and nutrient composition. Traditional forward genetics allows genetic mapping, while functional phenomics links phenes to plant performance. Based on these data, genotypes with contrasting phenotypes can be selected for smaller yet more intensive experiments to understand phene-environment interactions in depth. Simulation modeling is employed to understand the phenotypes and all stages of the pipeline feed back to ideotype and phenotyping platform development.

### Box 3. Phenetic hierarchies and phene integration

The ‘rocketship’ model is a useful visualization for understanding phenetic hierarchy and phene integration (detailed in York *et al.*, 2013). The phenes including root angle, total root number, photosynthesis rates, root cortical area, and exudation all influence plant functions such as nutrient acquisition, nutrient utilization, and the total carbon economy. Root number is a phene aggregate that decomposes into the more elemental phenes number of whorls and number of roots per whorl (York and Lynch, 2015). Similarly, cortical area is influenced by aerenchyma, cortical cell file number, and cortical cell size (Jaramillo *et al.*, 2013). The relations of lower order and greater order phenes constitute a phenetic hierarchy that acknowledges the uncertainty of knowing whether a proposed phene is actually elemental. At the same time, the contemplation of the elemental degree of a phene in relation to other phenes is a useful activity for ideotype development and design of phenotyping platforms. New classifications of phenes, their relations, and how they influence various plant function will benefit the utility of functional phenomics to deliver solutions. Phene integration determines how individual phenes interact to influence plant functions (York *et al.*, 2013). For example, in maize more aerenchyma decreases root segment respiration and therefore allows a more efficient use of photosynthate to construct longer roots, while reduced crown root number allows individual root axes to be allocated more photosynthate. Together, these two phene states allow greater nitrogen uptake, shoot mass, and yield. Understanding phenetic hierarchies and phene integration will allow greater gains to be made in ideotype breeding programs.

### Box 4. Recent progress in functional phenomics

#### High-throughput phenotyping of water use and plant size combined with linear modelling offers new insights to water use efficiency

A plant-to-sensor conveying platform was used to automatically measure water use via differences in pot weight and digital imaging to estimate biomass (Feldman *et al.*, 2017). Since total water use and total biomass were correlated, water use efficiency was explored using linear modelling to look at genotype-level deviations from the modelled relation. This research combined gains in understanding of plant physiology with genetic mapping, a powerful approach for functional phenomics.

#### Automated simulations of light interception using image-based plant architecture and biomass were used to calculate radiation use efficiency

Cabrera-Bosquet *et al.* (2016) combined a plant-to-sensor system measuring image-based shoot architecture with measurements of greenhouse light levels to simulate light interception and calculate radiation use efficiency. This approach show the power of combining simulation and experimental approaches.

#### Phenotyping leaf transpiration rate and phosphorus concentration linked chickpea water use to phosphorus uptake

Using 266 chickpea genotypes, Pang *et al.* (2018) used measures of phosphorus (P) concentration and shoot biomass to determine P-use efficiency and P-utilization efficiency. Using a subset of 100 lines contrasting for growth, they uncovered differences for photosynthetic rate and a positive correlation between transpiration and P acquisition. These results give an intriguing example of a non-intuitive relationship being uncovered through a functional phenomics approach, namely between mass flow and P uptake.

#### Reduced nighttime transpiration has been demonstrated to be an important breeding target in grapevine

Decreased transpiration at night was found to reduce water use without affecting growth in grapevine, representing a new avenue for increasing overall transpiration efficiency (Coupel-Ledru *et al.*, 2016). This system coupled conveyors, automated pot weighing and water for water use, and digital imaging for estimating biomass.

#### Root crown phenotyping linked root system architectural phenes to rooting depth and yield in wheat

In a wheat mapping population, Slack *et al.* (2018)found that root crown architectural phenes correlated to rooting depth, yield, and senescence phenology based on root crown phenotyping methodology described in York *et al.* (2018). Specifically, steeper root angles and more roots per shoot were associated with more root length density at 40 – 60 cm dpeth, which was, in turn, related to yield and canopy stay-green.

**Figure 1.**
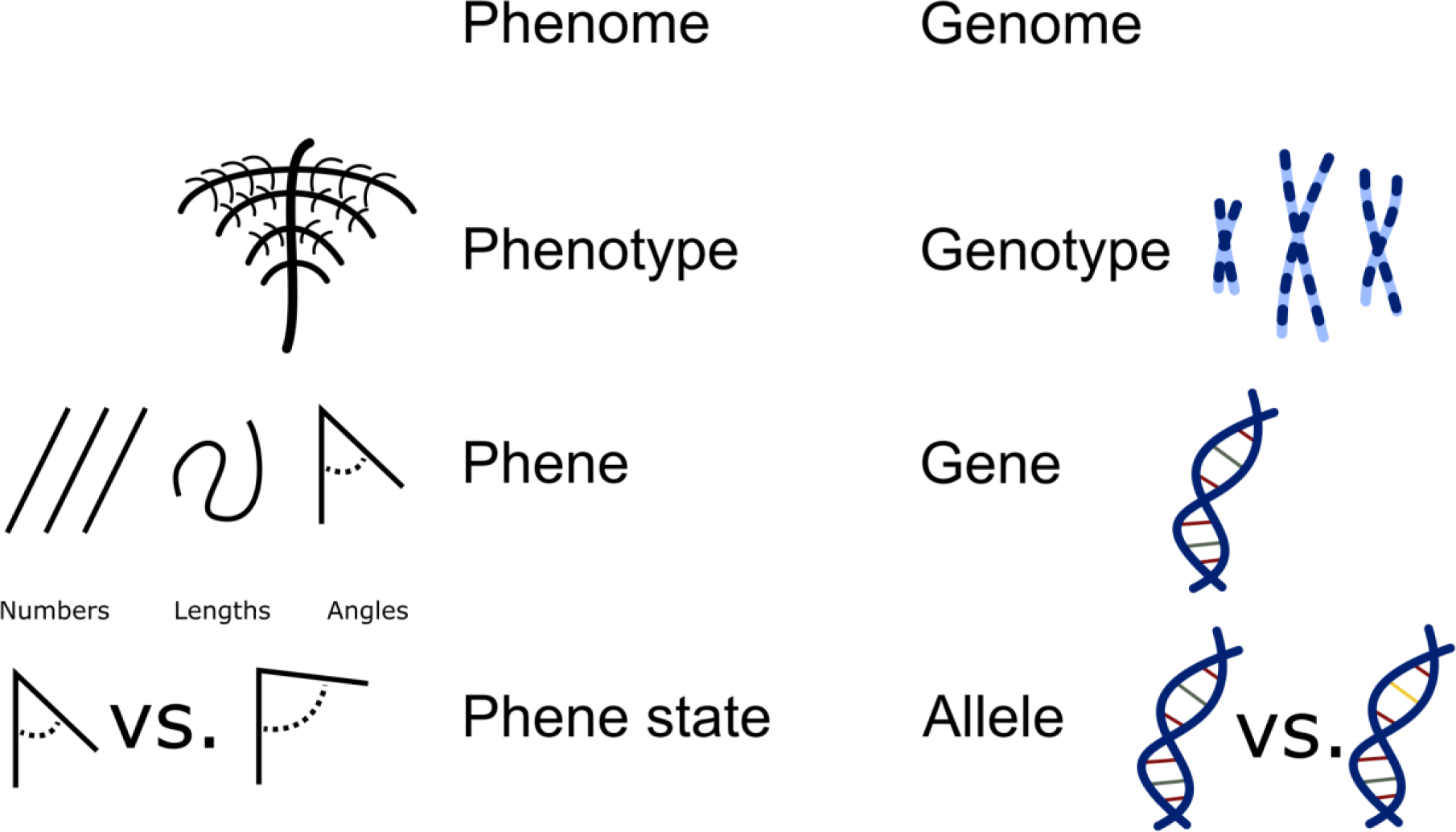
A 1:1 mapping of phenetic to genetic terms is proposed.

**Figure 2.**
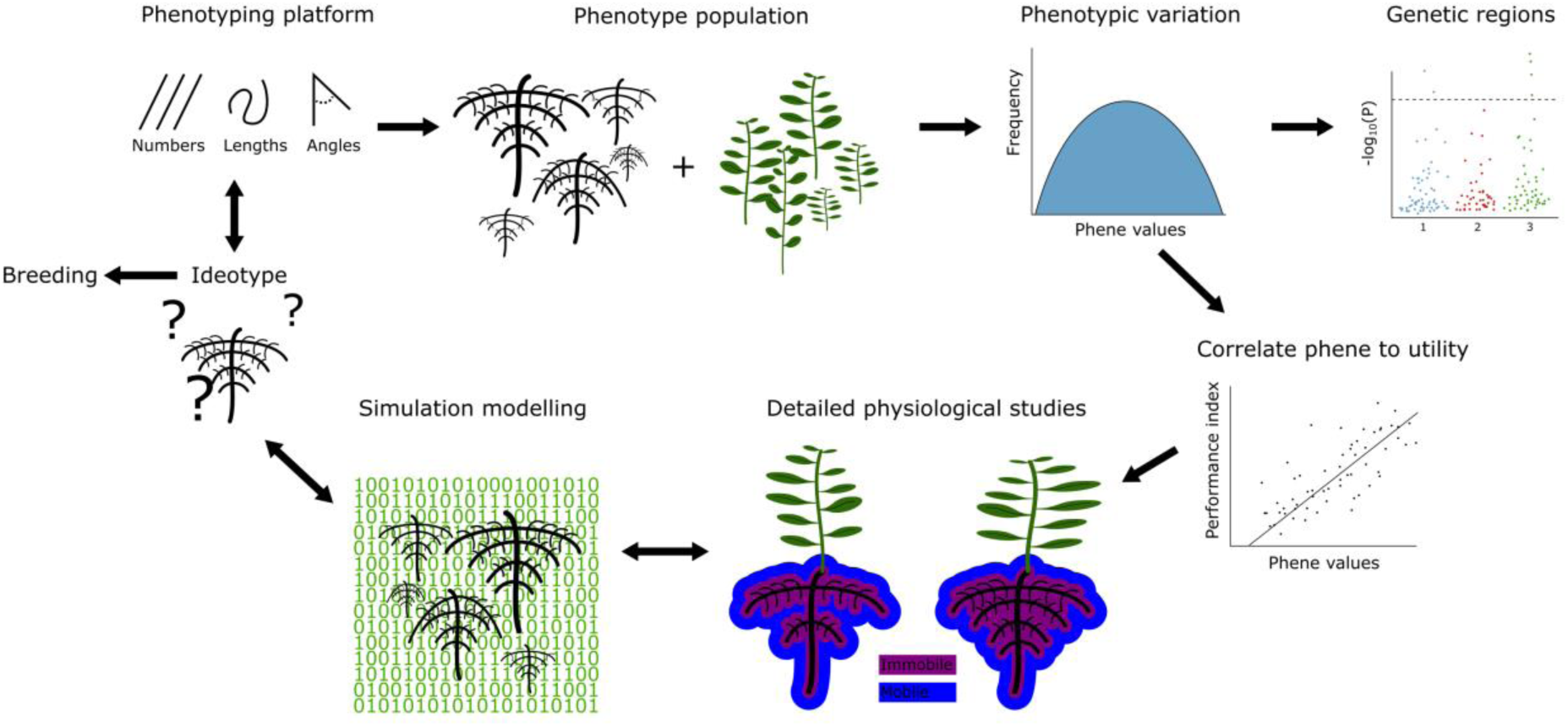
The functional phenomics pipeline is designed to increase fundamental knowledge of plant physiology by developing ideotypes, designing phenotyping platforms, phenotyping populations, studying phenotypic variation, correlating phenes to utility, physiological studies, and simulation modelling.

**Figure 3.**
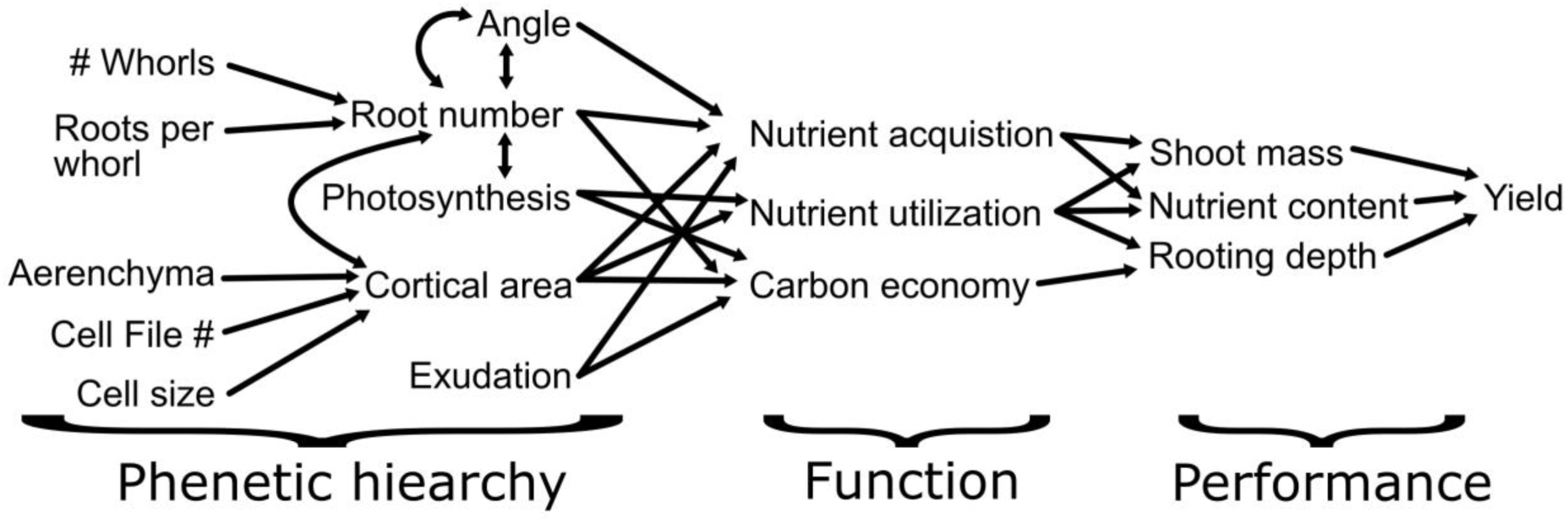
The ‘rocketship’ diagram demonstrates how the phenetic hierarchy leads to phene interactions that influence functional aspects and eventually performance metrics. Similar diagrams have been used by Arnold (1983), Violle *et al.* (2007), and York *et al.* (2013).

## Acknowledgements

I thank all my mentors who have influenced and prioritized my thinking on phenes and the importance of plant physiology for solving global challenges. I thank researchers from around the world for their hard work in this new field of functional phenomics and say carry on. I apologize for not being able to include all recent examples related to functional phenomics; exclusion does not imply anything about merit.

